# Collagen4 fluorescent protein knock-in lines reveal remodeling of cardiovascular basement membranes

**DOI:** 10.64898/2026.10.29.762927

**Authors:** Astha Tuladhar, Elizabeth J Wunderlich, Shawn A Slaymaker, Aishwarya Pawar, Maura McGrail, Pascal Lafontant, Jeffrey J Essner

## Abstract

Basement membranes (BMs) are specialized extracellular matrix structures that provide structural support and regulate cellular behavior. Although BM composition and organization are dynamically remodeled during development and tissue regeneration, the remodeling of individual BM components in the cardiovascular system (CVS) remain incompletely characterized. Here, we focused on Collagen4 (Col4), a major structural component of BMs, to investigate its remodeling during cardiovascular development and regeneration. To monitor the remodeling of Col4 *in vivo*, the Venus fluorescent tag was integrated in frame into the zebrafish endogenous *col4a1* and *col4a2* genes, enabling live confocal imaging of expression. Our data reveal dynamic patterns of Col4a1 and Col4a2 within the developing CVS, with deposition progressively increasing as vascular and cardiac structures develop and mature during both development and regeneration. In the regenerating heart, Col4a2 accumulated at the injury site during early stages of regeneration and became increasingly associated with the newly regenerated myocardium. Together, these findings demonstrate that Col4 deposition is dynamically reorganized across developmental and regenerative stages of CVS and describe new tools for following Col4 dynamics in vivo.

## Introduction

Basement membranes (BM) are a thin layer of specialized extracellular matrix (ECM) that forms an interface between cells and the surrounding tissue (Jayadev and Sherwood, 2017; Leclech et al., 2020). In addition to providing support to cells and tissues, BMs also help regulate signaling pathways that govern cell processes such as differentiation, migration, and proliferation (Randles et al., 2020; Schlie-Wolter et al., 2013; Sherwood, 2021). In cardiac tissues, BMs are present between endocardium and myocardium and between epicardium and myocardium (Boland et al., 2021; Jallerat and Feinberg, 2020). In the vasculature, BMs are present surrounding the blood vessels between the endothelial cells and pericytes (LeBleu et al., 2007; Leclech et al., 2020). In both the heart and vasculature, BMs are remodeled during development and regeneration as tissues expand and endothelial cells form new blood vessels during angiogenesis (Horne-Badovinac, 2020; Matsubayashi, 2022; Matsubayashi et al., 2020).

Collagen4 (Col4) is a major structural component of the BM, where it forms an interconnected network together with other BM components, including laminin, nidogen, and perlecan (Frantz et al., 2010; LeBleu et al., 2007; Leclech et al., 2020). Col4 consists of six distinct alpha chains, a1–a6, encoded by *COL4A1-COL4A6* in humans (Gelse, 2003). These six alpha chains are conserved among humans, mice, and zebrafish (LeBleu et al., 2023). Specifically, three alpha chains associate to form a triple-helical Col4 structure, with the a1a1a2 heterotrimer being predominant in cardiovascular BMs (Boland et al., 2021; Pöschl et al., 2004).

Col4 undergos dynamic remodeling during development and tissue morphogenesis. For example, live imaging using an eGFP-Col4a2 mouse model revealed dynamic changes in Col4a2 deposition and organization during hair follicle morphogenesis (Wuergezhen et al., 2024). However, relatively few studies have examined the spatiotemporal remodeling of Col4 specifically within the cardiovascular system (CVS). The distribution and organization of Col4 in the developing mouse ventricle was examined from embryonic day E12.5 to postnatal day P2, showing that Col4 deposition increased and became more prominent as the heart matured (Hanson et al., 2013). Proteomic analysis of the regenerating zebrafish heart further revealed temporal changes in ECM composition, including changes in Col4 (Garcia-Puig et al., 2019), but the spatial dynamics of Col4 deposition during cardiac regeneration was not evaluated. Additionally, the spatiotemporal dynamics of Col4 deposition during early cardiac development, vascular development, and regeneration remain largely unexplored due to a limited ability to image Col4 in vivo in living animals

Characterizing changes in Col4 deposition is particularly relevant given the dynamic nature of cardiovascular development and regeneration. During vascular development, blood vessels undergo remodeling as newly formed vessels sprout, branch, and remodel to establish a functional vascular network (Ellertsdóttir et al., 2010; Okuda and Hogan, 2020). Similarly, the developing heart undergoes progressive organization of the endocardium, myocardium, trabeculae, valves, and epicardium (Brown et al., 2016; Hallab et al., 2021). These dynamic changes require continuous adaptation of the surrounding extracellular environment (Lindsey and Borg, 2010; Sherwood, 2021). Tissue regeneration also requires remodeling of the ECM as damaged tissue is removed, and new cells populate the injury site (Garcia-Puig et al., 2019; Keane et al., 2018; Wang et al., 2023, 2024). Understanding how Col4 deposition is coordinated with these processes is crucial, especially as regenerative and tissue-engineering strategies increasingly utilize ECM-based scaffolds to recapitulate aspects of the native extracellular environment (Belviso et al., 2022, 2020; KC et al., 2019; Ott et al., 2008). Defining when and where Col4 is deposited and remodeled may therefore provide insights into ECM organization that could inform the design of regenerative and tissue-engineering approaches. To address this, we generated knock-in zebrafish reporter lines in which the Venus fluorescent protein was inserted into the endogenous *col4a1* and *col4a2* loci, enabling *in vivo* visualization of Col4 deposition. Using these reporter zebrafish lines, we analyzed the spatiotemporal dynamics of Col4 deposition during CVS development and regeneration. We demonstrate Col4 undergoes dynamic spatiotemporal deposition during development and regeneration, confirming a critical role for BM/ECM remodeling in during these processes and establishing new knock-in lines to follow BM dynamics.

## Results

### Generation of knock-in reporter zebrafish lines to analyze the spatiotemporal deposition of Col4

To map the spatiotemporal distribution of Col4 in the BM of the CVS, stable zebrafish lines were generated by inserting a Venus fluorescent protein coding sequence flanked by a short flexible linker on both ends into a CRISPR/Cas9 cut site in the N-terminus 7S domain of the *col4a1* and *col4a2* genes (Fig. 1A, B). This was achieved using GeneWeld CRISPR short homology-mediated targeted integration with a plasmid containing the linkers and Venus coding sequences as a template (Welker et al., 2021). The insertion site was selected in the 7S domain based on a previously established *eGFP-Col4a2* knock-in mouse model (Wuergezhen et al., 2024). Following injection, F0 embryos were screened for Venus fluorescence to identify embryos carrying insertions at the endogenous *col4a1* or *col4a2* loci. Screening of injected F0 embryos at the *col4a1* locus revealed moderate to extensive fluorescence in 18.5% (20/108) of the injected embryos. Similarly, screening at the *col4a2* locus identified fluorescence in 19.1% (13/68) of the injected embryos. Fluorescent F0 embryos were subsequently raised to adulthood and screened for germline transmission of the Venus tag. Germline transmission of Venus fluorescent protein was observed in 33.3% (2/6) of *col4a1* founders and 20% (1/5) of *col4a2* founders. Precise integration of the Venus was verified by sequencing the 5’ and 3’ junctions flanking the insertion site (Fig. S1, S2).

**Fig. 1.**
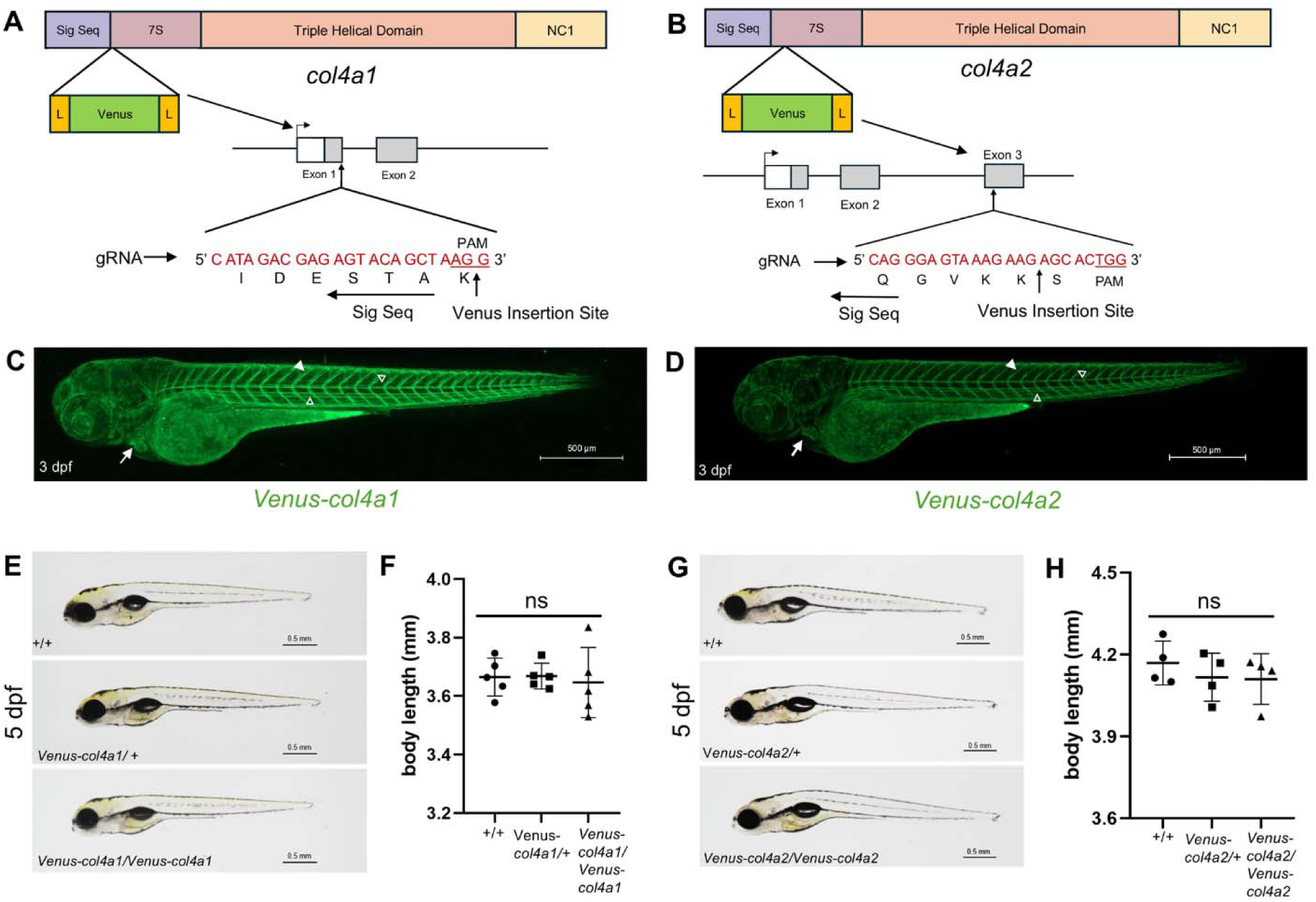
Generation of the Col4a1 and Col4a2 knock-in reporter lines. (A, B) Schematic illustration of *col4a1* (A) and *col4a2* (B) gene structures showing different domains of the gene: signal sequence (sig seq), N-terminus non-collagenous region (7S), triple helical domain, and C-terminus non-collagenous region (NC1), and the insertion site for Venus fluorescent protein. The Venus fluorescent protein flanked by short flexible linkers (L) and was inserted in-frame at the N-terminal region within the 7S domain of each gene. Gray indicates coding sequence and white the 5’ untranslated region. (C, D) Whole-body confocal imaging of *Venus-col4a1* and *Venus-col4a2* knock-in reporter lines showed localization of Col4a1 (C) and Col4a2 (D)to the myoseptum (arrowhead), vasculature (open arrowheads), and cardiac tissue (arrow). (E) Representative images of *Venus-col4a1/+* incross progeny at 5 dpf showing wild-type (+/+), heterozygous (*Venus-col4a1*/+), and homozygous (*Venus-col4a1*/*Venus-col4a1*) genotypes. (F) Quantification of body length at 5 dpf in *Venus-col4a1/+* incross progeny across the indicated genotypes (n=5). (G) Representative images of *Venus-col4a2/+* incross progeny at 5 dpf showing wild-type (+/+), heterozygous (*Venus-col4a2*/+), and homozygous (*Venus-col4a2*/*Venus-col4a2*) genotypes. (H) Quantification of body length at 5 dpf from *Venus-col4a2/+* incross progeny with the indicated genotypes (n=4). (C, D) scale bar = 500 μm, (E, G) scale bar = 0.5 mm. (F, H) One-way ANOVA (Tukey’s multiple comparisons test); ns, not significant; data are presented as mean ± SD.

The Tg(*Venus-col4a1*)*^is114^*and Tg(*Venus-col4a2*)*^is115^* knock-in reporter zebrafish lines exhibited indistinguishable expression patterns of Venus-Col4a1 and Venus-Col4a2 (Fig. 1C, D), consistent with the known assembly of Col4a1 and Col4a2 into a1a1a2 heterotrimers that form the triple-helical Col4 scaffold of BMs (Gelse, 2003) and the mRNA expression patterns for each gene (Flatman et al., 2025; Naylor et al., 2022; Sur et al., 2023). Consistent with the stoichiometry of the heterotrimers (Gelse, 2003), the *Venus-col4a1* line was brighter than the *Venus-col4a2* line. Analysis of *Venus-col4a1* and *Venus-col4a2* revealed prominent Col4 deposition within the myoseptum, cardiac tissue, and vasculature (Fig. 1C, D). The localization in the myoseptum, vasculature, and developing heart was consistent with the localization of Col4 in mice and humans (Jones et al., 2023; Kleppel et al., 1989; Steffensen et al., 2021; Wuergezhen et al., 2024). Homozygous knock-in zebrafish were viable and exhibited no overt morphological abnormalities during development (Fig. 1E-H). Body length measurements at 5 days post fertilization (dpf) were comparable between homozygous knock-in, heterozygous knock-in, and wild-type zebrafish (Fig. 1F, H), suggesting that the integration of the Venus tag does not interfere with the normal function of Col4a1 and Col4a2.

### *Venus-col4a1* and *Venus-col4a2* revealed dynamic spatiotemporal distribution of Col4 during development of vasculature

To visualize endothelial cells for colocalization with Venus-Col4a1 and Venus-Col4a2, the Tg(*fli1b-gal;14XUAS-RFP)* or Tg(*kdrl:mCherry)* vascular endothelial red fluorescent reporter lines were used (Balciuniene and Balciunas, 2013; Bertrand et al., 2010). Live confocal imaging at 1 dpf revealed that Col4a1 and Col4a2 were detected in the myoseptum, along the abluminal side of the dorsal aorta ventral to the hypocord, but no detectable expression was observed around the developing intersegmental vessels (ISVs) (Fig. 2A, B). A single optical section of Venus-Col4a2 shows two layers of deposition of Col4a2 along the abluminal side of the dorsal aorta and adjacent to the hypochord at 1 dpf (Fig. S3A, arrowhead and arrow). By 2 dpf, Col4a1 and Col4a2 expression was evident around the dorsal aorta, ISVs, and dorsal longitudinal anastomotic vessel (DLAV) on the abluminal side of the patent vessel (Fig. 2C, D). Imaging of the caudal vein plexus from 2-3 dpf using *Venus-col4a2* zebrafish line further revealed that Col4a2 deposition in the caudal vein closely mirrors alterations in vascular architecture (Fig. S3B, C). A higher-magnification image at 3 dpf demonstrated the abluminal localization of Col4a2 along the dorsal aorta and ISVs (Fig. S4). Both Venus-Col4a1 and Venus-Col4a2 deposition were progressively more prominent around the developing vasculature at successive stages during vascular development and maturation (Fig. 2).

**Fig. 2.**
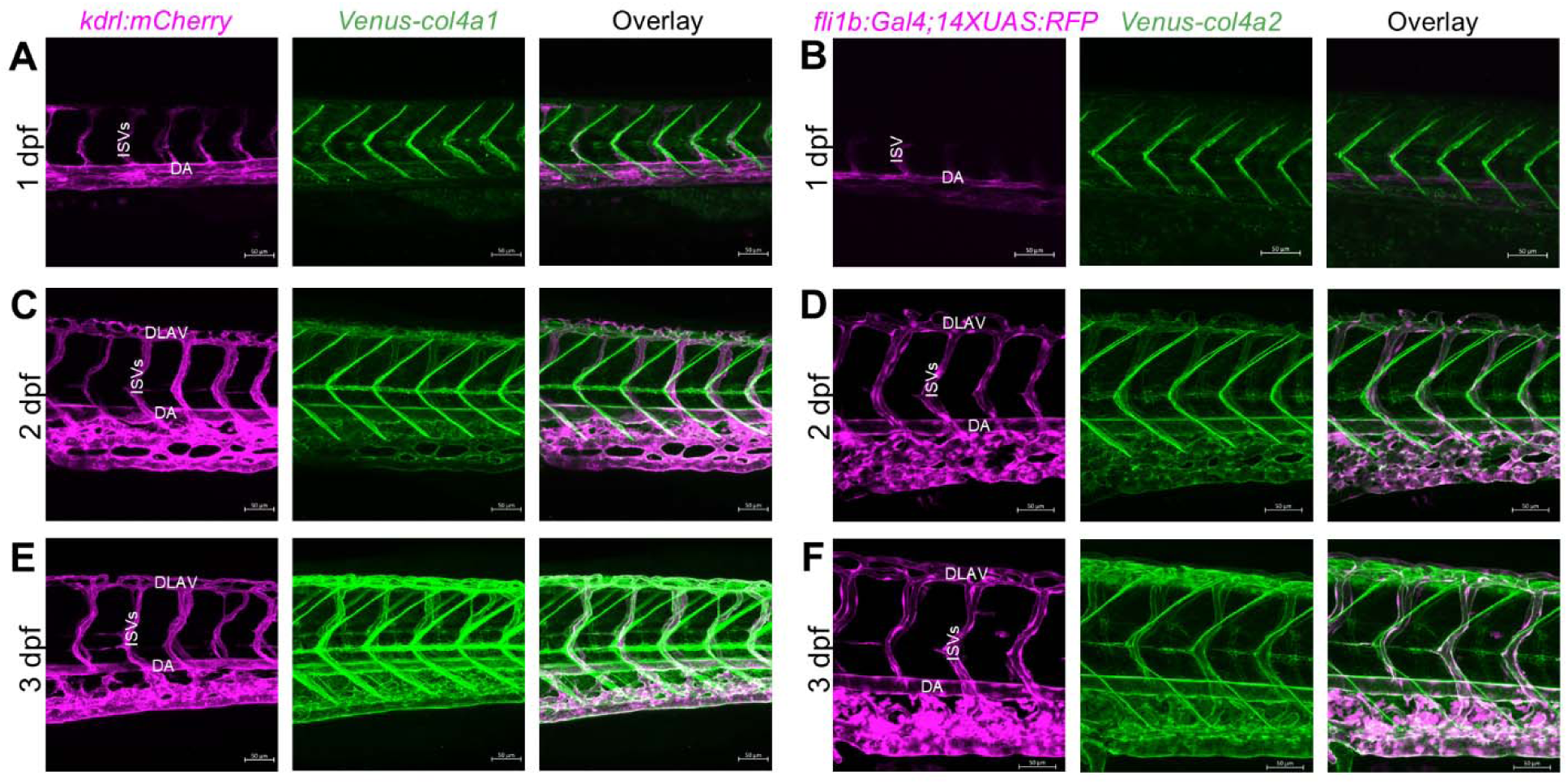
Dynamic remodeling of Col4a1 and Col4a2 during vascular development. (A-F) Live confocal imaging of the zebrafish embryo showing the spatiotemporal deposition of Col4a1 and Col4a2 in relation to the developing vasculature from 1-3 dpf. Images show the dorsal aorta (DA), intersegmental blood vessels (ISVs), and dorsal longitudinal anastomotic vessel (DLAV). Scale bar = 50 μm.

Since Col4a2 and Col4a1 deposition at the ISVs was first detected at 1 dpf, when blood flow is established, we next asked whether this deposition is influenced by blood flow. To test this, we quantified Venus-Col4a2 fluorescence in *silent heart/tnnt2* mutants which lack a beating heart and control siblings at 2 dpf using the Tg(*tnnt2-rfp*) line (Clark et al., 2011). Homozygous *tnnt2^-/-^* mutants exhibited a significant increase in Venus-Col4a2 fluorescence at the dorsal aorta compared with heterozygous *tnnt2^-/+^*mutants (Fig. 3A, B). In contrast, Venus-Col4a2 fluorescence surrounding the ISVs was reduced in *tnnt2^-/-^*mutants relative to control (Fig. 3A, C). These opposing effects in the silent heart mutants likely indicate blood flow differentially regulates Col4a2 deposition around the dorsal aorta and ISVs.

**Fig. 3.**
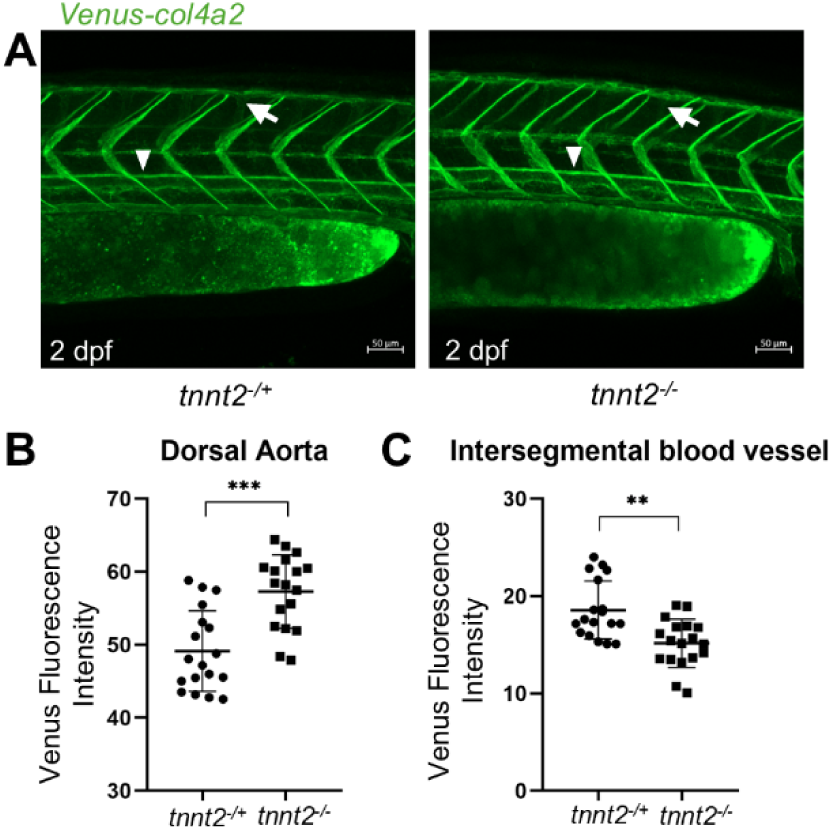
Effects of loss of blood flow on Col4a2 deposition. (A) zebrafish embryos at 2 dpf showing Venus-Col4a2 deposition in heterozygous (with a heartbeat) and homozygous (without a heartbeat) *silent heart* mutants around the dorsal aorta (arrowhead) and ISVs (arrow). (B, C) Quantification of fluorescence intensity of Venus-Col4a2 in the dorsal aorta (B) and intersegmental blood vessels (C) in the *silent heart* mutants. Scale bar = 50 μm. Student’s t-test; **, P < 0.01; ***, P < 0.001.

### *Venus-col4a1* and *Venus-col4a2* revealed deposition of Col4 during development of cardiac tissue

We next examined the spatiotemporal deposition of Col4a1 and Col4a2 during cardiac development. To visualize the colocalization of Venus-Col4a1 and Venus-Col4a2 near the developing cardiomyocytes, *Tg*(*tnnt2-rfp*) was used. Col4a1 was detectable as early as 1 dpf along the inner surface of the myocardium (Fig. 4A, open arrowhead). In contrast, Col4a2 was first detected at 2 dpf along the inner surface of the myocardium (Fig. 4B, open arrowhead). Differences in gene expression and subsequent fluorescence intensity between the two reporter lines may explain the earlier detection of Col4a1. To further spatially define the localization of Col4, *Tg(kdrl:mCherry)* was used to detect the co-localization of Venus-Col4a2 in relation to the endocardium. Col4a2 was deposited on the outer surface of the endocardium (Fig. S5), suggesting that Col4a2 is deposited within the BM between the endocardium and myocardium. By 3 dpf, another layer of Col4a1 deposition was observed along the outer surface of the cardiomyocytes (Fig. 4E, arrowhead), whereas an outer layer of Col4a2 deposition was detected by 4 dpf (Fig. 4F, arrowhead). This deposition coincided with the development of the epicardial layer covering the myocardium (Boezio et al., 2022). Expression of both Col4a1 and Col4a2 were more prominent at later developmental stages (Fig. 4), indicating a progressive accumulation during heart development. Further, Col4a1 and Col4a2 deposition was also detected in other cardiac structures. *Venus-col4a2* optical sections from 2–5 dpf showed Col4a2 deposition in developing trabeculae (arrow) and the outflow tract (arrowhead) (Fig. S6).

**Fig 4.**
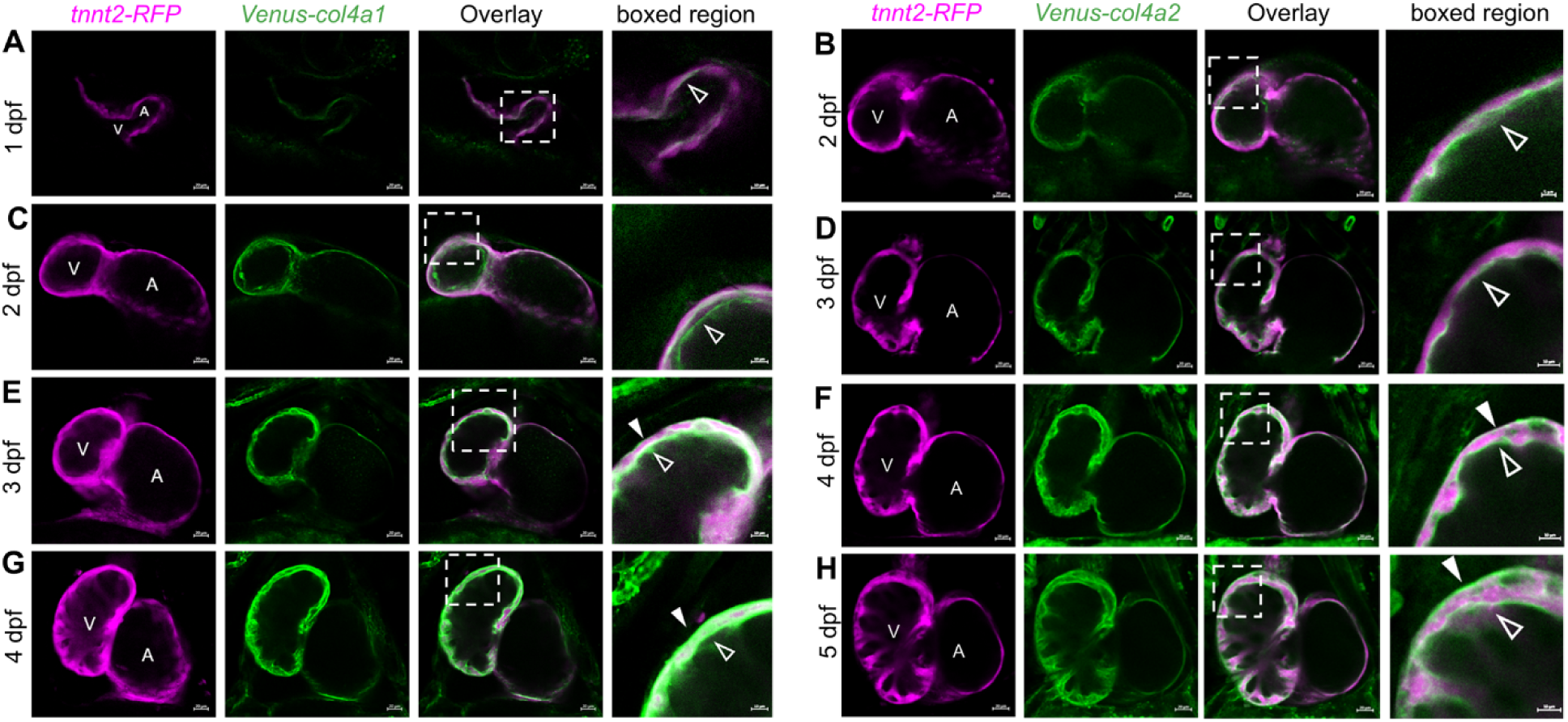
Deposition and remodeling of Col4a1 and Col4a2 during heart development. Confocal imaging of heart development, showing the spatiotemporal distribution of Col4a1 at 1 dpf (A), 2 dpf (C), 3 dpf (E), and 4 dpf (G) and Col4a2 at 2 dpf (B), 3 dpf (D), 4 dpf (F), and 5 dpf (H) in relation to the developing cardiomyocytes in single Z-sections to ventricle (V) and atrium (A). (A-H) Scale bar = 20 μm; boxed regions have scale bars of 5 μm (B) and 10 μm (A, C-H).

### *Venus-col4a2* revealed dynamic spatiotemporal distribution of Col4a2 during regeneration of vasculature

Having established the deposition of Col4a2 and Col4a1 during cardiovascular development, we next examined whether Col4 undergoes dynamic remodeling during cardiovascular regeneration in zebrafish. Since the *Venus-col4a1* and *Venus-col4a2* lines showed similar overall patterns of localization during development, we focused on the *Venus-col4a2* line for the subsequent regeneration studies. To investigate the spatiotemporal dynamics of Col4 during vascular regeneration, we used larval and adult zebrafish caudal fin amputation models. The caudal fin fold in larval zebrafish was amputated at the position indicated in Fig. S7A which severs the connection of the dorsal aortal and caudal vein. Live imaging of the amputated larval fin fold revealed that Col4a2 first accumulated within the regenerating tissue and became increasingly associated with regenerating vasculature as regeneration proceeded (Fig. S7). At 1-day post-amputation (dpa), we detected Col4a2 starting to accumulate in the regenerating tissue (Fig. S7B, arrow). Time-lapse imaging of the fin immediately after the amputation further demonstrated Col4a2 starting to populate the regenerating tissue and being remodeled as the dorsal aorta and caudal vein reconnect to establish a circulatory loop (Video S1). By 2 dpa, as regeneration progresses and blood vessels begin to invade the regenerating tissue, little to no Col4a2 was detected (Fig. S7C). As vascular regeneration progresses, we detected more Col4a2 deposition in the mature vasculature (arrow) as compared to the immature vasculature (arrowhead) (Fig. S7D, E). Analysis of adult fin regeneration revealed a similar pattern of Col4a2 remodeling (Fig. 5). At 1 dpa, we detected Col4a2 expression in the blastema (Fig. 5A, B, open arrowhead). By 3 dpa, the regenerating fin had extended, and new blood vessels were evident within the regenerating tissue, with Col4a2 expression observed in the regenerating vasculature (Fig. 5A). Col4a2 deposition was more prominent around mature vasculature (arrow) compared to immature vasculature (arrowhead), as shown in the higher-magnification image of the 3 dpa regenerating vasculature (Fig. 5C). At 7 dpa, the regenerating fin and vasculature had extended further, and Col4a2 deposition showed a similar pattern to that observed at 3 dpa with higher deposition around mature vessels more proximal to the site of amputation (arrow) (Fig. 5A).

**Fig 5.**
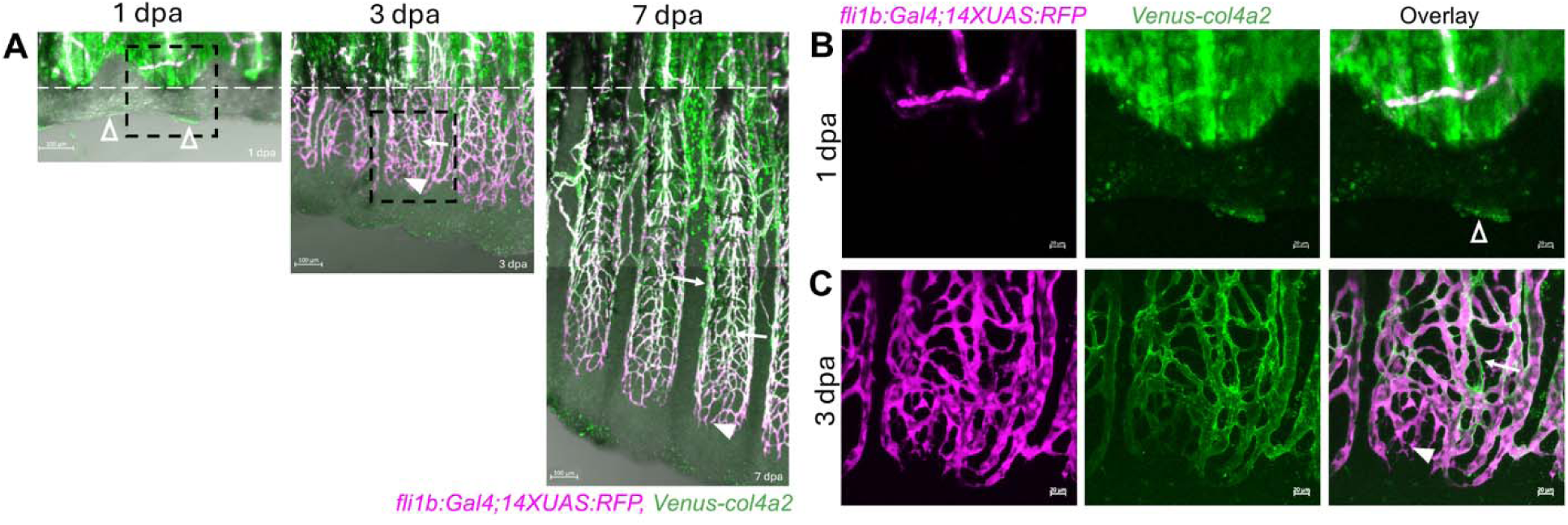
Deposition and remodeling of Col4a2 during vascular regeneration in adult zebrafish. (A) Confocal imaging of adult zebrafish showing the spatiotemporal deposition of Col4a2 during vascular regeneration at 1 dpa, 3 dpa, and 7 dpa. The white dotted line indicates the site of fin amputation. Col4a2 deposition is detected within the blastema at 1 dpa (open arrowhead). As vasculature regeneration progresses to 3 and 7 dpa, Col4a2 deposition becomes more prominent around the mature regenerating vasculature (arrow) than around the immature regenerating vasculature (arrowhead). (B, C) Higher magnification images of the boxed region in A at 1 dpa (B) and at 3 dpa (C). (A) Scale bar = 100 μm, (B, C) scale bar = 20 μm.

### *Venus-col4a2* revealed a dynamic spatiotemporal distribution of Col4a2 during regeneration of cardiac tissue

We next examined the spatiotemporal distribution of Col4a2 during heart regeneration using heart resection in double transgenic *tnnt2-rfp;Venus-col4a2* adults. Cross-sections of regenerating hearts were immunolabeled with antibodies against RFP and GFP to visualize cardiomyocytes and Col4-Venus, respectively. In the adult heart, Col4a2 was localized around the outer layer surrounding the compact myocardium (outer muscular layer of the ventricular wall) (Fig. 6A arrowhead) and underlying the myocardial trabeculae (Fig. 6A arrow). Col4a2 was deposited at the site of injury during the early stage of regeneration, as demonstrated by its localization at the site of amputation at 3 dpa (Fig. 6B arrow). By 7 dpa, during active regeneration (Lien et al., 2012; Major and Poss, 2007), the Col4a2 deposition is seen within the regenerating tissue (Fig. 6C arrow) as well as surrounding the regenerating tissue (Fig. 6C arrowhead), with Col4a2 preceding the population of the regenerating cardiomyocytes in the new tissue. By 14 dpa, when regenerated myocardium progressively replaced the injured tissue (Lien et al., 2012; Major and Poss, 2007), we observed a reduction in Col4a2 deposition in the regenerating tissue (Fig. 6D arrowhead). At this stage, Col4a2 was also detected surrounding the regenerated cardiomyocytes (Fig. 6D arrow), indicating progressive re-establishment of BM architecture as the myocardium regenerated. By 30 dpa, new cardiomyocytes had replaced the damaged tissue and Col4a2 deposition appeared comparable to the unamputated heart, localizing around the compact myocardium (Fig. 6E arrowhead) and underlying the trabecular myocardium (Fig. 6E arrow). Together, these findings demonstrate that Col4a2 undergoes remodeling during cardiac regeneration.

**Fig. 6.**
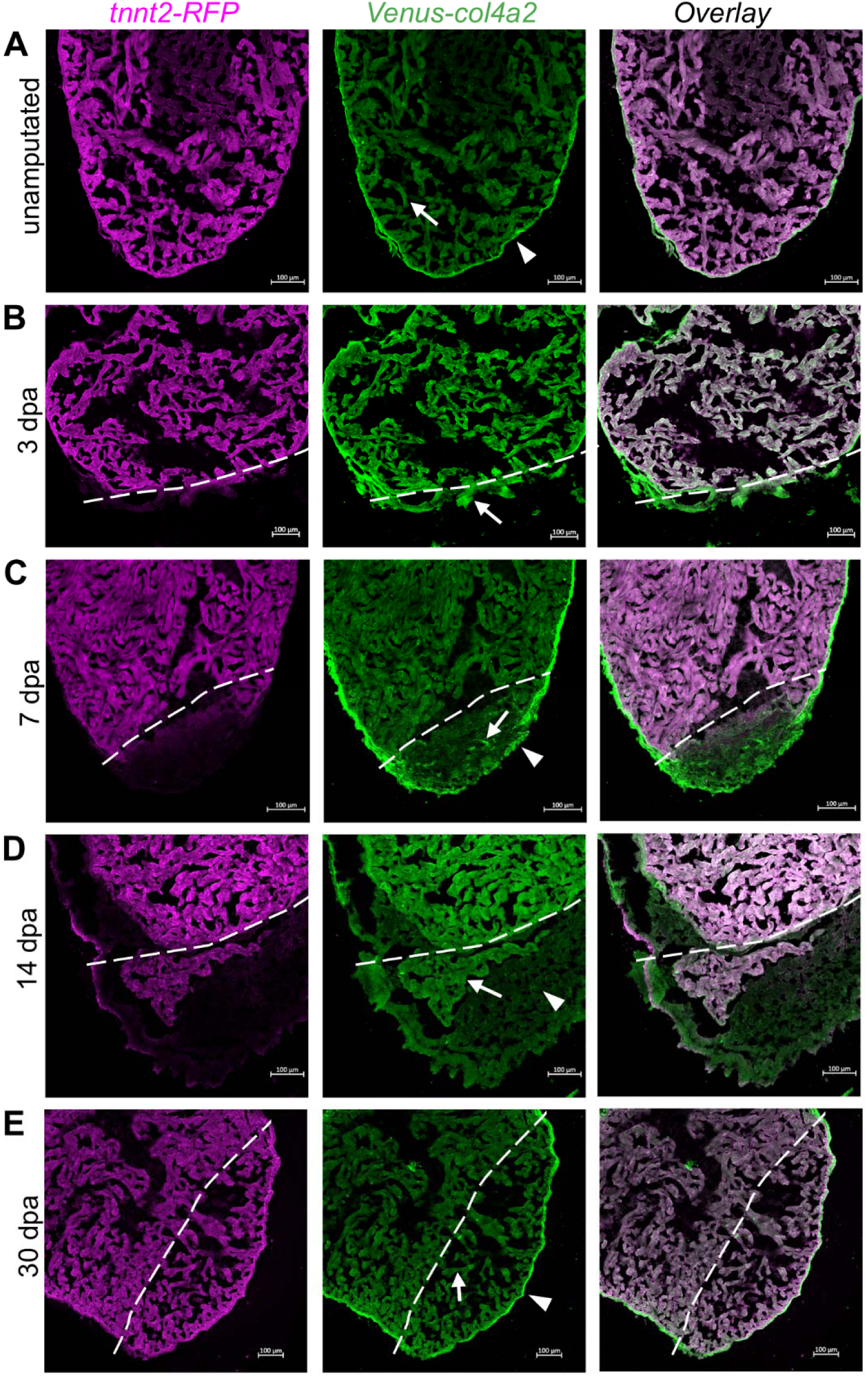
Deposition and remodeling of Col4a2 during heart regeneration in adult zebrafish. (A) Representative confocal image of an uninjured heart showing Col4a2 deposition around the compact myocardium (arrowhead) and in association with the trabecular myocardium (arrow). (B–E) Representative confocal images during heart regeneration showing the spatiotemporal deposition of Col4a2 relative to regenerating tnnt2-RFP+ cardiomyocytes. The white dotted line indicates the site of injury. (B) Representative confocal image at 3 dpa showing Col4a2 deposition at the injury site (arrow). (C) Col4a2 deposition within the regenerating tissue (arrow) and surrounding the wound region (arrowhead) at 7 dpa. (D) Col4a2 deposition was enriched in association with the newly regenerated cardiomyocytes (arrow) compared with the surrounding regenerating tissue (arrowhead) at 14 dpa. (E) Col4a2 deposition around the compact myocardium (arrowhead) and in association with the trabecular myocardium (arrow) at 30 dpa, resembling the distribution observed in the uninjured heart in A. Scale bar = 100 μm.

## Discussion

We describe *Venus-col4a1* and *Venus-col4a2* knock-in zebrafish reporter lines to visualize Col4 *in vivo* dynamics. We characterized the spatiotemporal deposition of both Col4a1 and Col4a2 within cardiovascular BMs during development, and the remodeling of Col4a2 during fin and cardiovascular regeneration. Our findings reveal dynamic patterns of Col4 deposition associated with vascular and cardiac development, as well as dynamic remodeling during tissue regeneration.

During vascular development, Col4a1 and Col4a2 deposition closely followed the establishment and maturation of the vasculature. Col4a1 and Col4a2 deposition was first detected around the dorsal aorta at 1 dpf and around ISVs by 2 dpf, with fluorescence becoming progressively more prominent as the vessels matured. Our findings suggest that loss of cardiac contraction and subsequent blood flow reduced Col4a2 accumulation in the ISVs but increased accumulation around the dorsal aorta. These differences may be related to differences in developmental timing and the distinct mechanisms by which the dorsal aorta and ISV form and mature. The dorsal aorta is established early as part of the primary axial vasculature through vasculogenesis, whereas ISVs form later through angiogenic sprouting from the dorsal aorta (Lawson and Weinstein, 2002; Poole and Coffin, 1989). Further, during angiogenic sprouting, endothelial tip cells actively remodel and proteolytically degrade the surrounding extracellular matrix, including the vascular BM, to facilitate endothelial migration and invasion (Rowe and Weiss, 2008; Senger and Davis, 2011). Following sprout formation, BM components assemble and stabilize the newly established vessels (Senger and Davis, 2011; Stratman et al., 2009). Thus, the differences in deposition of Col4a1 and Col4a2 in dorsal aorta and ISVs may reflect differences in the extracellular environment associated with actively developing versus more established vessels. Consistent with this idea, Col4a2 deposition during vascular regeneration was more prominent around the mature vasculature than around newly regenerating vessels.

During cardiac development, Col4 deposition progressively appeared at distinct tissue interfaces as cardiac structures established and matured. Col4a1 was first detected at the myocardium–endocardium interface by 1 dpf, whereas Col4a2 was first detected at 2 dpf. By 3 dpf, we observed an additional layer of Col4a1 along the outer surface of the myocardium, while Col4a2 became detectable in this region by 4 dpf. Since Col4a1 and Col4a2 assemble into the heterotrimeric network (a1a1a2) (Khoshnoodi et al., 2008), the differences in the timing of detectable fluorescence may therefore reflect differences in signal intensity and the limits of detection rather than a complete absence of Col4a2. The distinct deposition of Col4 seems to be associated with the organization and maturation of cardiac tissue rather than its initial formation. Deposition of Col4 becomes detectable between the myocardium and endocardium after the formation of the heart tube. Similarly, the deposition of Col4 in the outer layer of the myocardium coincides with the establishment of the epicardium (Serluca, 2008). Consistent with our observation, previous work done in developing mouse hearts also showed a progressive increase in its deposition during cardiac development (Hanson et al., 2013). However, Hanson et al. reported that Col4 was initially detected in association with the epicardium and myocardium, followed by more prominent localization to the endocardium.

During cardiac regeneration, we can detect Col4a2 at the site of injury at 3 dpa, and the deposition of Col4a2 seemed to increase by 7 dpa around the regenerating tissue, with the expression of Col4a2 preceding the population of the regenerating tissue with cardiomyocytes. Later, by 14 dpa, as the new regenerating myocardium replaces the regenerating area, the Col4a2 deposition is seen in association with the myocardium as opposed to in the regenerating tissue. This observation is consistent with proteomics analysis in the regenerating heart that showed an increase at 7 dpa (Garcia-Puig et al., 2019). Our imaging analysis further provides spatial information that was not captured by the proteomic analysis. The early deposition of Col4a2 at the injury site may reflect active ECM remodeling during regeneration and its incorporation into the extracellular environment supporting tissue repair (Garcia-Puig et al., 2019; Mercer et al., 2013). As new myocardium forms, the later association of Col4a2 with the regenerated tissue may reflect the re-establishment of the BM around the newly formed myocardium as the heart regains its normal organization.

This study defines the spatiotemporal distribution of Col4 in the CVS. The *Venus-col4a1* and *Venus-col4a2* lines can be used to directly address the in vivo dynamics of Col4a1 and Col4a2; however, the functional consequences of Col4 remodeling remain unclear. Previous studies in mice as well as in zebrafish have shown that Col4 contributes to tissue integrity and influences cell–matrix interactions and signaling (Boland et al., 2025; Favor et al., 2007; Flatman et al., 2025; Paradisi et al., 2026; Pöschl et al., 2004; Stratman et al., 2017). Further investigation of the functional role of Col4a1 and Col4a2 at different developmental and regenerative stages will be important for understanding the biological significance of the distinct spatiotemporal patterns. Col4 represents only one component of the BM. Our observations of distinct patterns of endogenous Col4 deposition support that studying the remodeling of other BM components will be important for determining how the BM as a whole is remodeled during cardiovascular development and regeneration.

## Materials and Methods

### Approval of animal experiments and zebrafish maintenance

All experiments described were approved by the Iowa State University IACUC protocol 25-143. Zebrafish (*Danio rerio*) were raised and maintained on an Aquaneering aquaculture system at 26 °C on a 14 hour light/10 hour dark cycle. The wild-type (WIK) zebrafish used for the experiments were obtained from Zebrafish International Resource Center.

### Transgenic zebrafish lines

The previously described transgenic lines *Tg(kdrl:HsHRAS-mCherry)^s896^* (Bertrand et al., 2010); Tg(*fli1b^tpl50Gt^*) (Craig et al., 2015); Tg(*miniTol2 <14XUAS:mRFP, gCry:GFP>*)^tpl2^ (Balciuniene and Balciunas, 2013); and Tg(*tnnt2-rfp^mn0031Gt^)* (Clark et al., 2011) were used. The Tg(*fli1b^tpl50Gt^*) line used in this study was modified by FLP recombinase to remove the UAS-GFP sequence. For this study the transgenic lines Tg(*fli1b^tpl50Gt^*), Tg(*kdrl:HsHRAS-mCherry)^s896^*, Tg(*miniTol2 <14XUAS:mRFP, gCry:GFP>*)^tpl2^, and *tnnt2-rfp^mn0031Gt^* are referred to as *fli1b:Gal4, kdrl:mCherry, 14XUAS:RFP,* and *tnnt2-rfp,* respectively.

### Isolation of *Venus-col4a1 and Venus-col4a2* knock-in reporter zebrafish lines

The Tg(*Venus-col4a1*)*^is114^* and Tg(*Venus-col4a2*)*^is115^* knock-in lines were generated using the GeneWeld CRISPR/Cas9-mediated targeted integration method, as previously described (Wierson et al., 2020). Guide RNAs targeting the N-terminal 7S domain of *col4a1* (5′-CATAGACGAGAGTACAGCTA-3′) and *col4a2* (5′-CAGGGAGTAAAGAAGAGCAC-3′) were used to direct targeted integration of Venus into the genome. The plasmids containing homology arms for insertion of linker-Venus-linker into the *col4a1* and *col4a2* loci were constructed by HiFi Assembly (NEBuilder® HiFi DNA Assembly, New England Biolabs, Ipswich, MA, USA), using the pGFuse vector (Welker et al., 2021) as PCR template. Primers used for HiFi cloning are listed in Table 1. Three PCR fragments were amplified and assembled using NEBuilder® HiFi DNA Assembly. Primer pairs used for plasmid construction were *Venus-col4a1* F1/*Venus*-R1, *Venus*-F2/*Venus-col4a1* R2, and *Venus-col4a1* F3/*Venus-col4a1* R3 for the *Venus-col4a1* construct, and *Venus-col4a2* F1/*Venus*-R1, *Venus*-F2/*Venus-col4a1* R2, and *Venus-col4a1* F3/*Venus-col4a1* R3 for the *Venus-col4a2* construct.

**Table 1.**
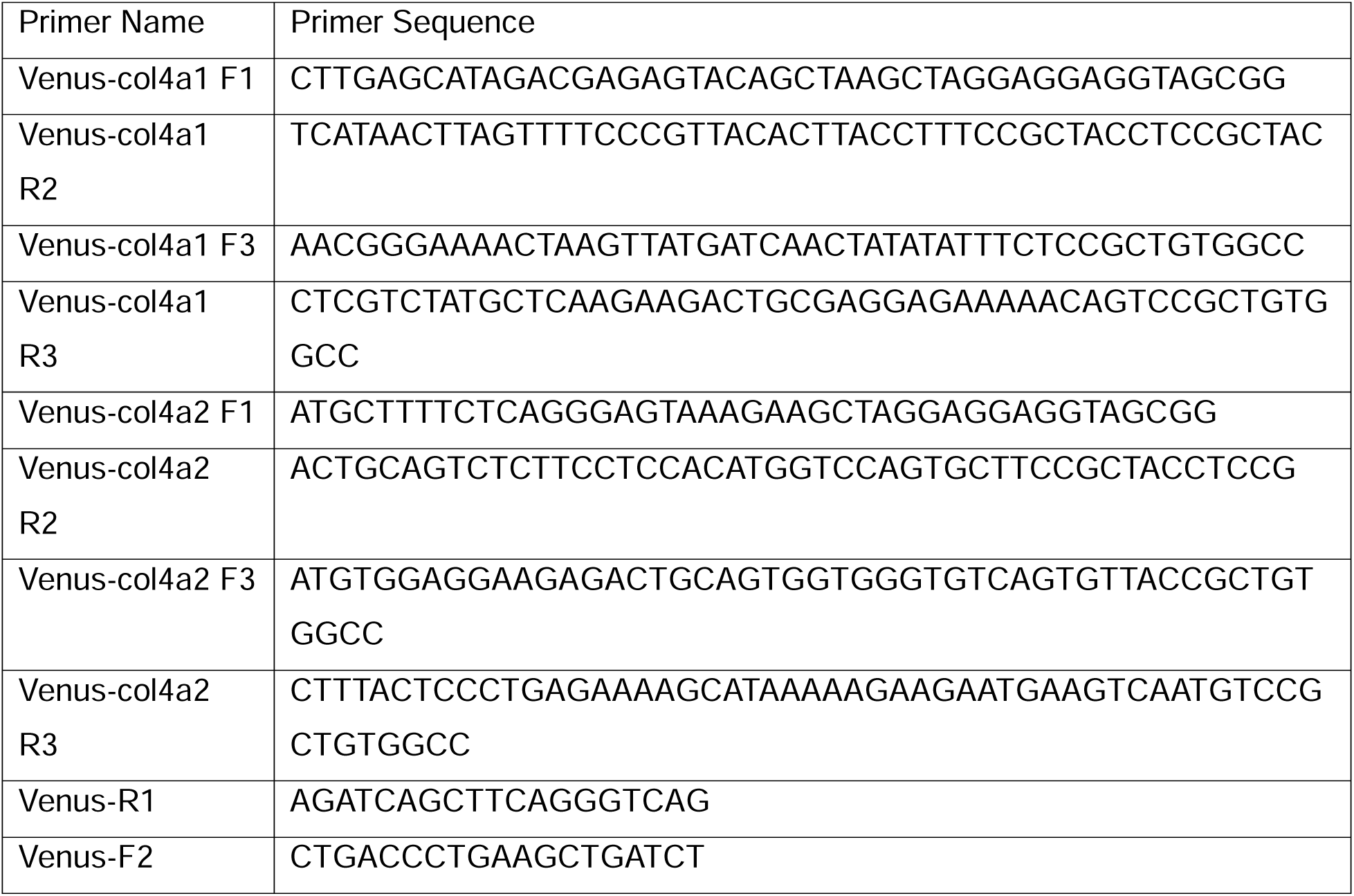
Primers used for HiFi cloning of templates for targeted integration.

### Genotyping

Genomic DNA was extracted from zebrafish embryos or adult caudal fin by digesting in 50 mM NaOH at 95°C for 30 min. The resulting DNA was used as the template for PCR-based genotyping. Additionally, to verify the 5′ integration junction of the *Venus-col4a2* knock-in allele, total RNA was first extracted using the RNeasy Mini Kit (QIAGEN, Hilden, Germany) and used as the template for RT-PCR performed using the Luna® One-Step RT-PCR Kit (New England Biolabs, Ipswich, MA, USA). Primer sequences used for genotyping are listed in Table 2.

**Table 2.**
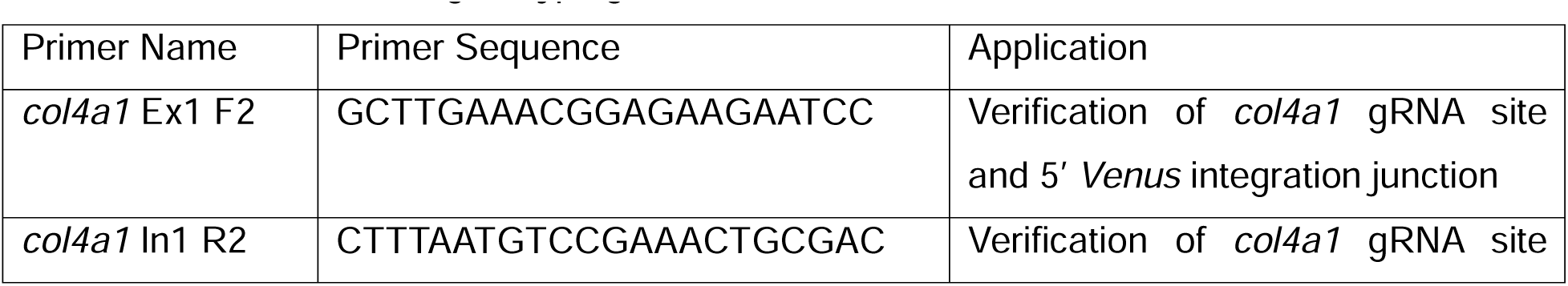
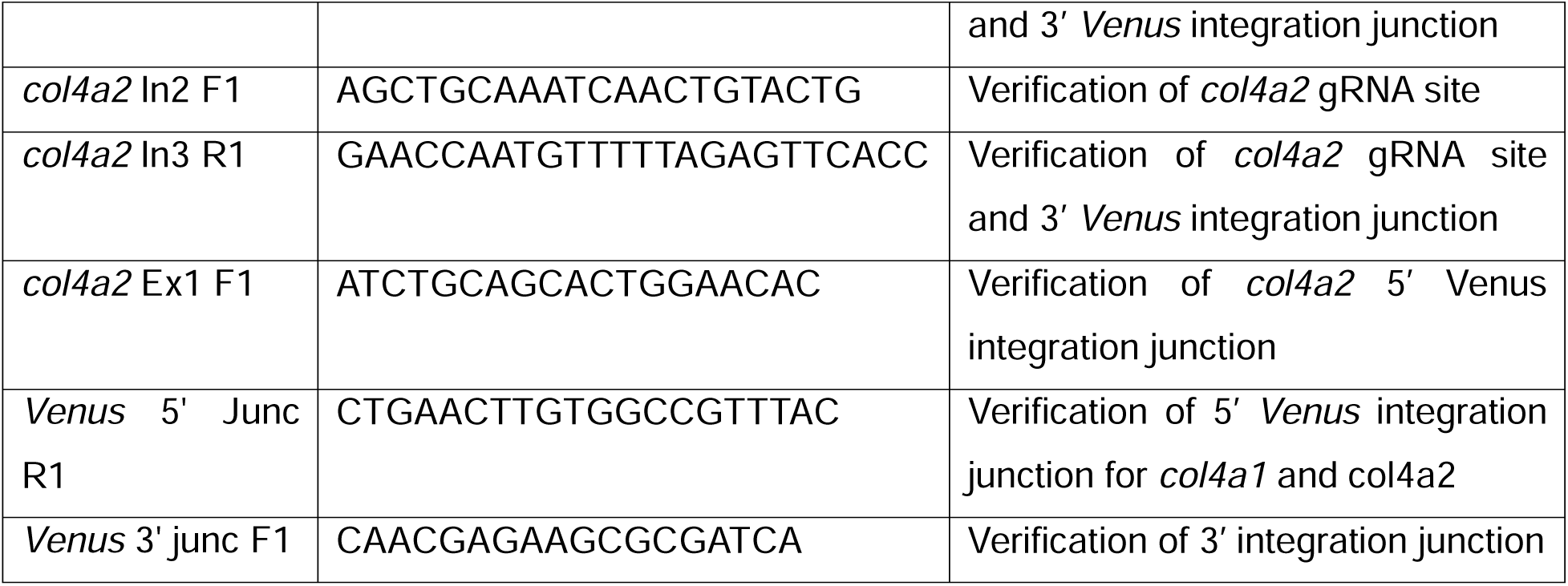
Primers used for genotyping the *Venus-col4a1* and *Venus-col4a2* alleles.

### Zebrafish Vascular Regeneration Assay

For larval vascular regeneration assays, 3 dpf larvae were anesthetized in 0.02% tricaine, and the caudal fin fold was amputated at the position indicated in Fig. S7A using fine iridectomy scissors. For the adult vascular regeneration assay, 3-6-month-old zebrafish were first anesthetized with 0.016% tricaine, and approximately half of the caudal fin was amputated with scissors. Following amputation, fish were returned to fresh system water and maintained under standard husbandry conditions until the indicated time points for imaging.

### Heart Resection

Heart resection was performed in adult zebrafish (3-6 months old) as previously described (Poss et al., 2002). Briefly, adult zebrafish were anesthetized in 0.016% tricaine and positioned ventral side up in a moist, slotted sponge. A small incision was made through the ventral body wall to expose the heart, and approximately 20% of the ventricular apex was surgically resected using fine iridectomy scissors. Following surgery, the fish were immediately returned to fresh system water for recovery and maintained under standard husbandry conditions. Hearts were collected at the indicated time points after amputation and processed for cryosectioning and immunostaining.

### Tissue processing and immunostaining

Adult zebrafish hearts were collected and fixed overnight at 4°C in 4% paraformaldehyde (PFA) prepared in phosphate-buffered saline (PBS). Following fixation, the hearts were washed with PBS and cryoprotected by incubation in 30% sucrose in PBS at 4°C for 48 h. The tissues were then embedded in OCT compound and stored at −80°C until cryosectioning. The frozen tissues were sectioned using a Thermo Scientific HM 550 cryostat. Sections were cut at a thickness of 12 μm. The tissue sections were mounted onto positively charged glass slides (Superfrost Plus Gold, Fisher Scientific) and stored at −20°C until further processing for immunostaining.

For immunostaining, the sections were washed three times for 5 min each in PBS containing 0.1% Triton X-100 (PBST) and blocked for 1 h at room temperature in blocking solution consisting of 10% lamb serum and 1% bovine serum albumin (BSA) prepared in PBST. The sections were then incubated overnight at 4°C with the primary antibodies: chicken anti-GFP (Aves Labs, GFP-1020; 1:300) and rabbit anti-RFP (Takara bio, Living Colors® DsRed Polyclonal Antibody, 632496; 1:300) were used at the indicated dilutions.

Following primary antibody incubation, the sections were washed three times for 10 min each with PBST and incubated with the appropriate fluorophore-conjugated secondary antibodies for 2 hours at room temperature in the dark. Secondary antibodies were used at a dilution of 1:500 for goat anti-rabbit Alexa Fluor 594 (Invitrogen, A-11012) and goat anti-chicken Alexa Fluor 488 (Invitrogen, A-11039). The sections were subsequently washed three times for 5 min each with PBST, mounted with 3-4 drops of Fluoro-Gel mounting medium containing DAPI (Electron Microscopy Sciences, 17985-50), and coverslipped. The slides were stored at 4°C until imaging.

### Imaging and Image Processing

For live imaging, zebrafish embryos were treated with 0.003% 1-phenyl-2-thiourea (PTU) (Sigma, P7629) before 24 hpf, anesthetized with 0.02% tricaine, and mounted in 1.5% low-melting-point agarose. For imaging of the larval heart, 0.12% tricaine was used to arrest cardiac contractions during image acquisition. For imaging of adult hearts, 3–4 sections centered in the mid-ventricular region of each heart were analyzed. Confocal images were acquired using a Zeiss LSM800 laser scanning microscope. Confocal images were processed and analyzed using Zen lite software (version 3.12).

### Statistics

Statistical analyses were performed using GraphPad Prism. Data are presented as mean ± standard deviation (SD). Student’s t-test was used for comparison between two groups. Comparisons between three groups were performed using one-way ANOVA followed by Tukey’s multiple comparisons test. A P value < 0.05 was considered statistically significant.

## Acknowledgements

We would like to thank Dr. Fang Liu (Iowa State University) for her guidance and training in experimental procedures.

## Competing Interests

JJE and MM have competing interests with Immusoft Inc., LifEngine and LifEngine Animal Health. All other authors do not have competing interests.

## Funding

This work was supported by the National Institutes of Health NIH ORIP R24OD020166, R24OD036201 (MM, JJE) and NSF 2533961 (JJE).

## Data and Resource Availability

The zebrafish lines and vectors generated in this study are available upon request (JJE).

## AI Statement

AI was used to correct grammar.

## Supplemental Figure Legends

**Fig. S1.**
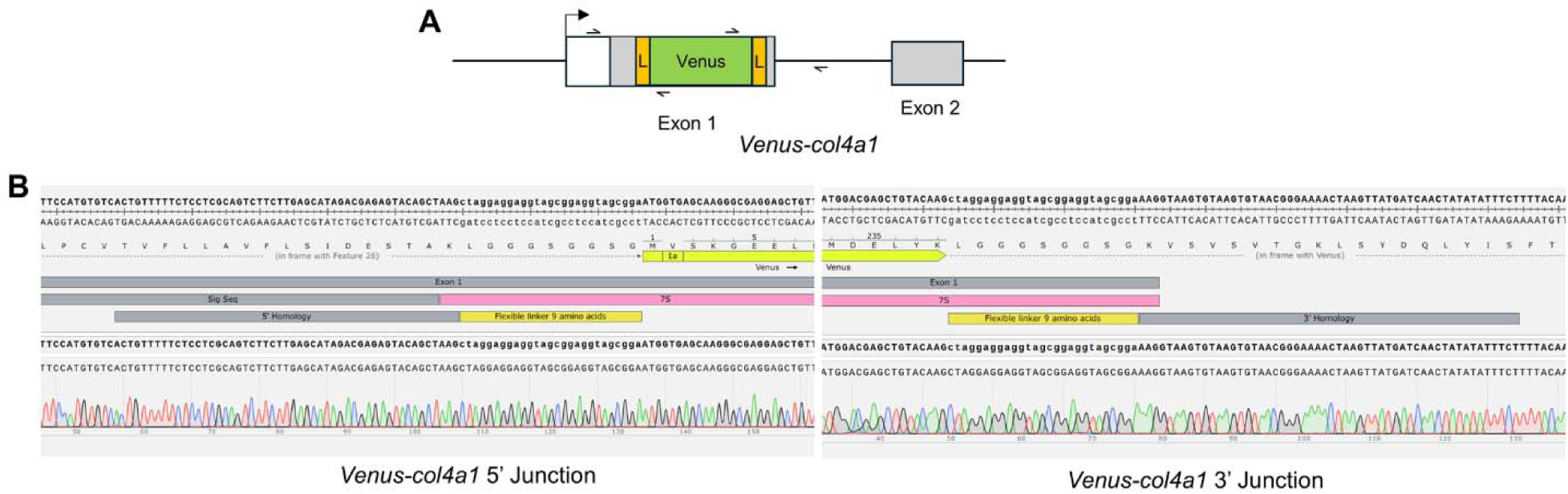
Junction verification of *Venus-col4a1* reporter zebrafish line. (A) Schematic of the *Venus-col4a1* locus showing primer binding sites used to verify the 5′ and 3′ integration junctions. (B) DNA sequences of the 5′ and 3′ integration junctions confirming the insertion of *Venus* at the *col4a1* locus. Gray indicates coding sequence and white the 5’ untranslated region. L-yellow boxes are the linker sequences.

**Fig. S2.**
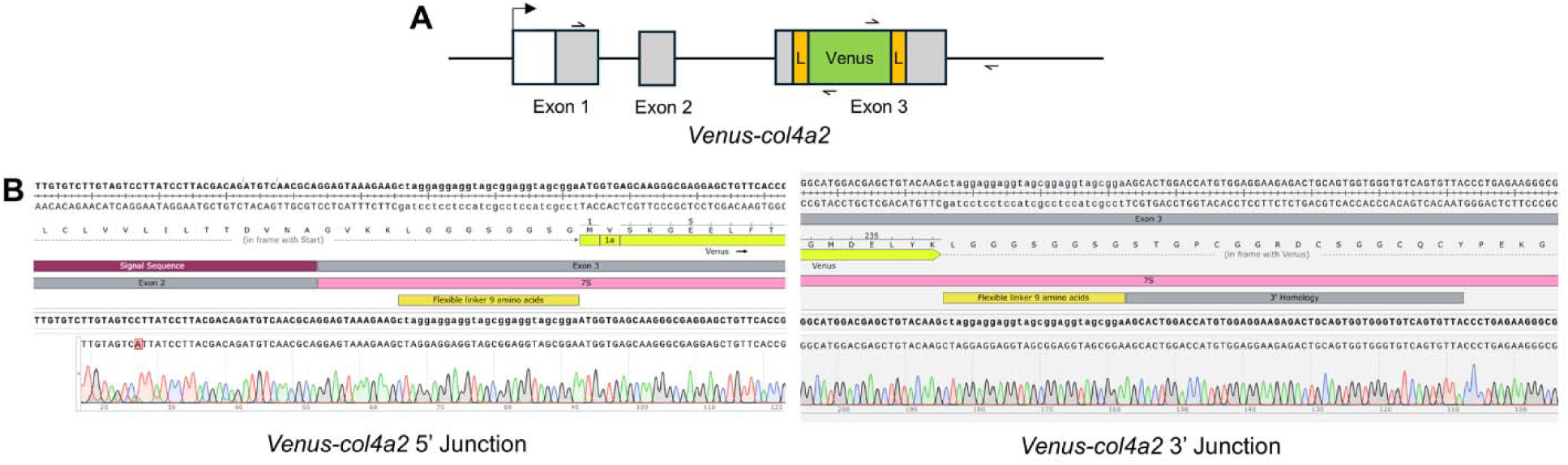
Junction verification of *Venus-col4a2* reporter zebrafish line. (A) Schematic of the *Venus-col4a2* locus showing primer binding sites used to verify the 5′ and 3′ integration junctions. (B) DNA sequences of the 5′ and 3′ integration junctions confirming the insertion of *Venus* at the *col4a2* locus. Gray indicates coding sequence and white the 5’ untranslated region. L-yellow boxes are the linker sequences.

**Fig. S3.**
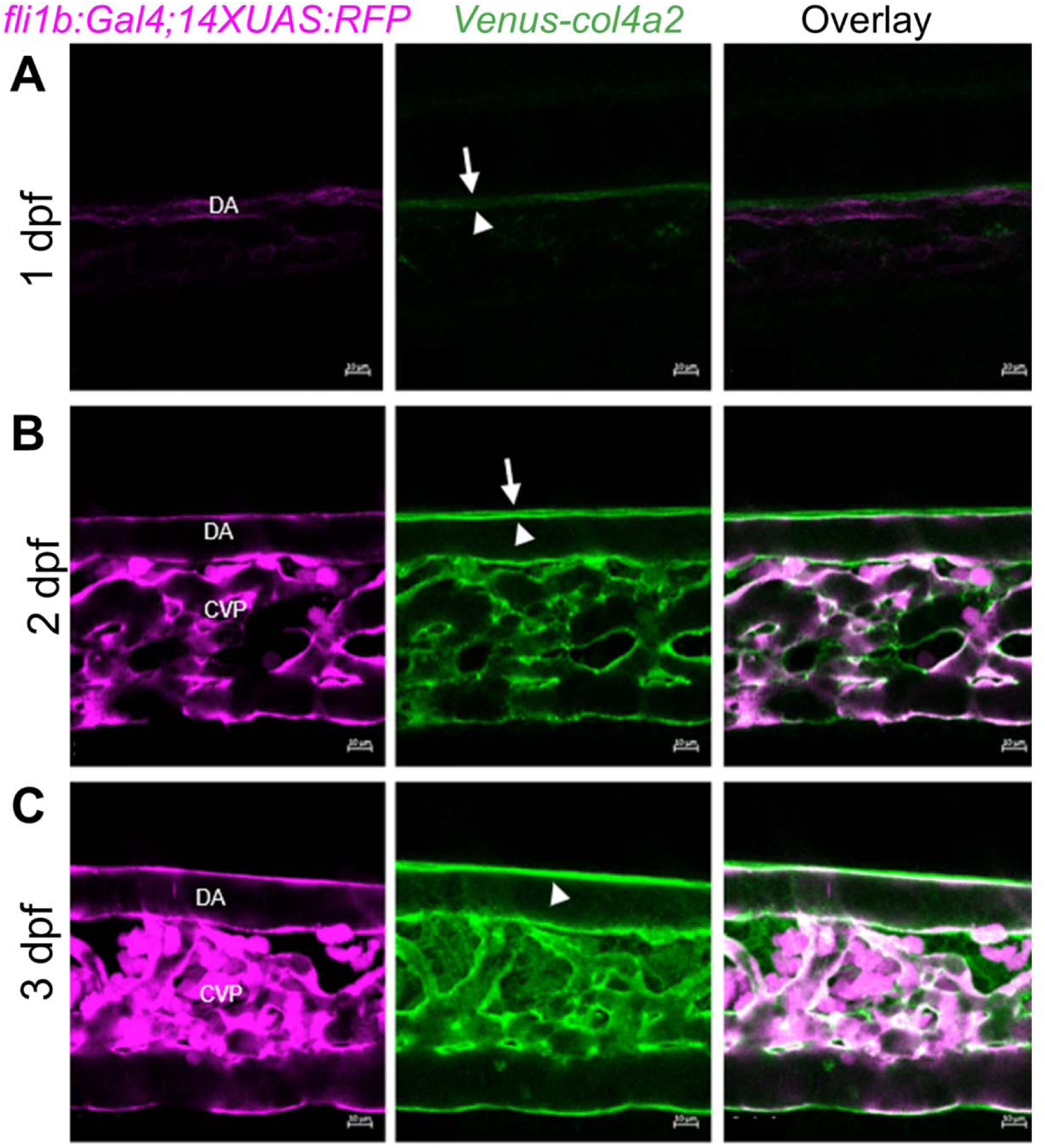
Single confocal sections showing remodeling of Col4a2 during vascular development. (A-C) Confocal image showing deposition of Col4a2 at 1 dpf (A), 2 dpf (B), and 3 dpf (C) in relation to hypochord (arrowhead), dorsal aorta (DA), and caudal vein plexus (CVP). Scale bar = 10 μm.

**Fig. S4.**
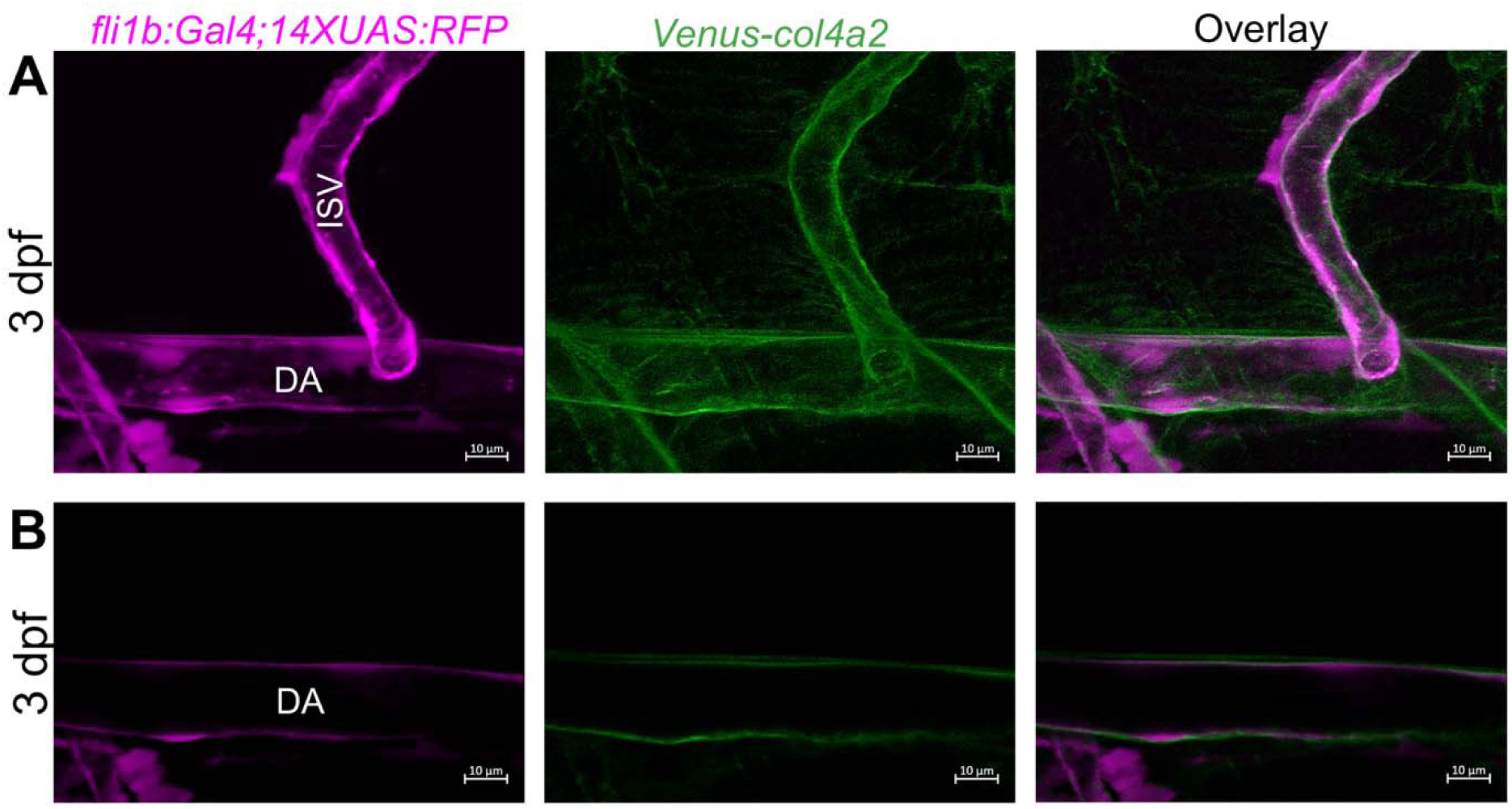
Deposition of Venus-Col4a2 around the vasculature at 3 dpf. (A) Confocal image showing deposition of Venus-Col4a2 on the abluminal side of the dorsal aorta (DA) and ISVs. (B) Single z-slice image showing the deposition of Venus-Col4a2 around the dorsal aorta. Scale bar = 10 μm.

**Fig. S5.**
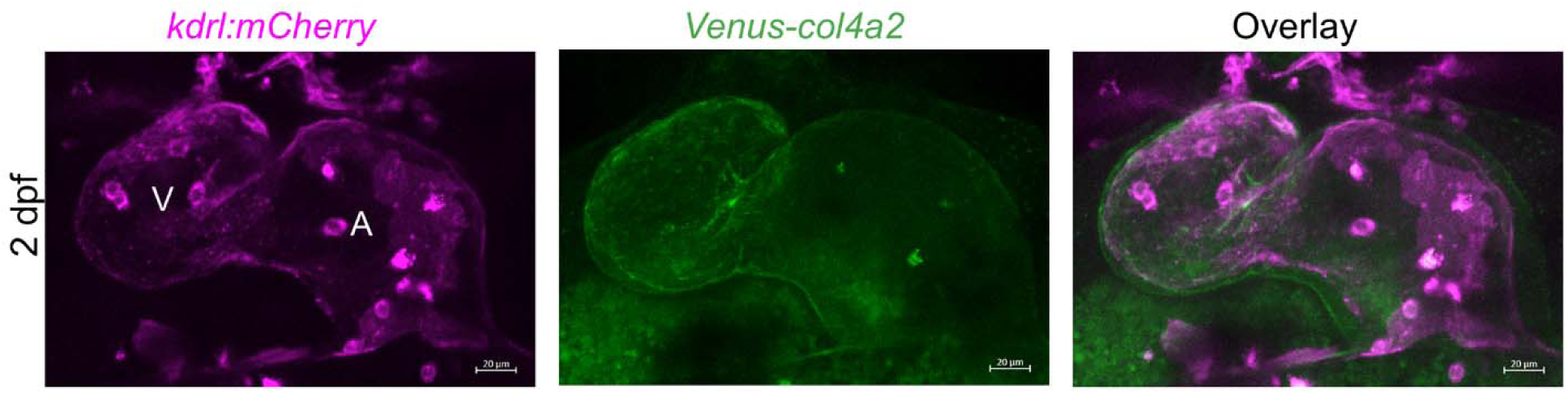
Deposition of Venus-Col4a2 in cardiac tissue with respect to the endocardium. (A) 2 dpf zebrafish heart showing localization of Venus-Col4a2 adjacent to the endocardium. V – ventricle, A – atrium. Scale bar = 20 μm.

**Fig. S6.**
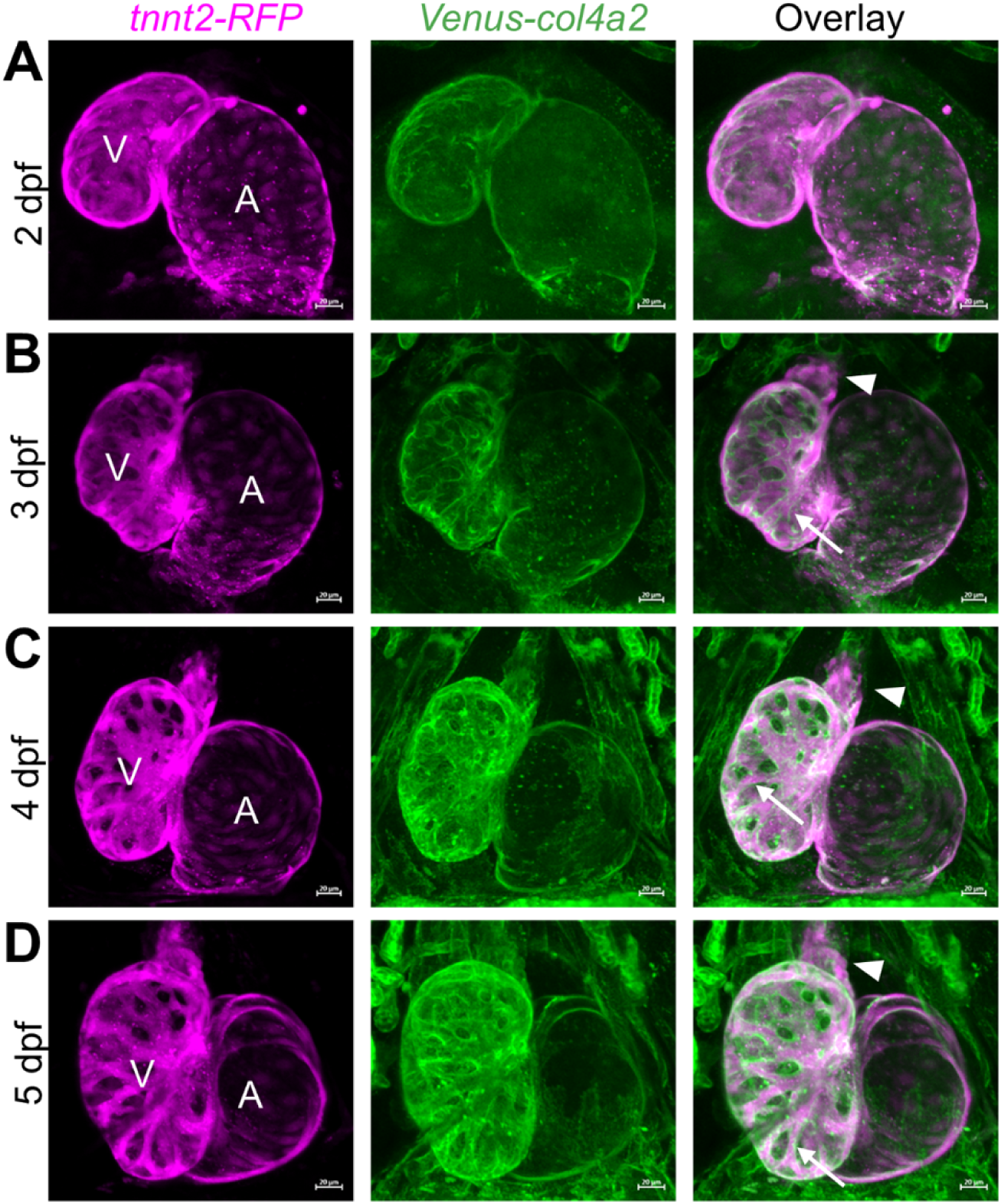
Dynamic remodeling of Col4a2 during cardiac development. (A-D) Confocal Z-stack imaging of *Venus-col4a2* showing the spatiotemporal distribution of Col4a2 at 2 dpf (A), 3 dpf (B), 4 dpf (C), and 5 dpf (D) in relation to the developing cardiomyocytes (tnnt2-rfp+ cells). Col4a2 deposition in the outflow tract is shown by the arrowhead and surrounding the trabeculae by the arrow. Scale bar = 20 μm.

**S7.**
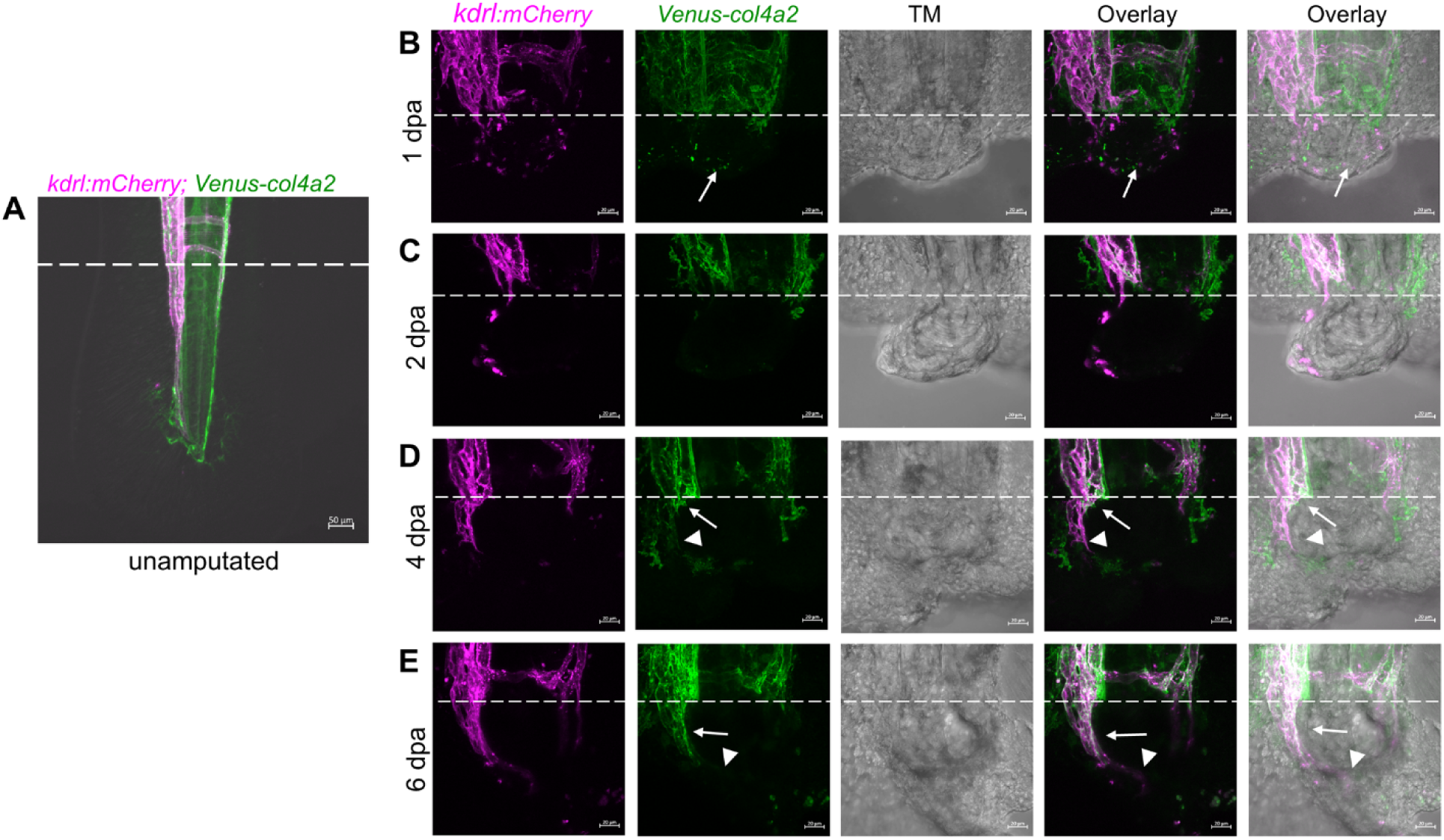
Col4a2 deposition during vascular regeneration in zebrafish larvae. (A) Confocal image showing the site of fin fold amputation (white dotted line) in a zebrafish larva. (B–E) Live imaging of zebrafish larvae showing the spatiotemporal deposition of Col4a2 during vascular regeneration at 1 dpa (B), 2 dpa (C), 4 dpa (D), and 6 dpa (E). The white dotted line indicates the site of amputation. (B) Col4a2 deposition is detected within the regenerating tissue associated with the *kdrl:mcherry+* endothelial cells at 1 dpa (arrow). (D, E) At 4 and 6 dpa, Col4a2 deposition is more prominent around mature regenerating vasculature (arrow) than around immature regenerating vasculature (arrowhead). (A) Scale bar = 50 μm, (B-E) Scale bar = 20 μm.

**Video S1.** Time-lapse confocal imaging of Tg(*kdrl:mCherry; Venus-col4a2*) zebrafish embryo immediately after fin amputation, showing Col4a2 (green) remodeling relative to vasculature (magenta) during fin regeneration. Reconnection of the dorsal aorta and caudal vein results in remodeling of the Venus-col4a2+ matrix. The movie was captured over 8 hours.

